# Functional differentiation in the language network revealed by lesion-symptom mapping

**DOI:** 10.1101/2020.07.17.209262

**Authors:** William Matchin, Alexandra Basilakos, Dirk-Bart den Ouden, Brielle C. Stark, Gregory Hickok, Julius Fridriksson

## Abstract

Theories of language organization in the brain commonly posit that different regions underlie distinct linguistic mechanisms. However, such theories have been criticized on the grounds that many neuroimaging studies of language processing find similar effects across regions. Moreover, condition by region interaction effects, which provide the strongest evidence of functional differentiation between regions, have rarely been offered in support of these theories. Here we address this by using lesion-symptom mapping in three large, partially-overlapping groups of aphasia patients with left hemisphere brain damage due to stroke (N=121, N=92, N= 218). We identified multiple measure by region interaction effects, associating damage to the posterior middle temporal gyrus with syntactic comprehension deficits, damage to posterior inferior frontal gyrus with expressive agrammatism, and damage to inferior angular gyrus with semantic category word fluency deficits. Our results are inconsistent with recent hypotheses that regions of the language network are undifferentiated with respect to high-level linguistic processing.

## 1 Introduction

Language is a multifaceted system consisting of interacting components. Beyond the phonetic and phonological levels, it consists of a set of lexical items (roughly words and morphemes), with associated conceptual representations, hierarchical syntactic structures, and complex semantic interpretation of these elements. Many authors have sought to associate these distinct aspects of language with different perisylvian brain regions thought to underlie language, often (but not always) supported primarily by neuroimaging data (Bornkessel-Schlesewsky & Schlesewsky, 2013; Friederici, 2017; Hagoort, 2014; Tyler & Marslen-Wilson, 2008; cf. Matchin & Hickok, 2020). However, the fact that a given neuroimaging study may find significant activations in some regions and not in others does not prove a functional distinction among them. In any given study, statistical power may be stronger in some regions rather than others.

Therefore, region by condition interaction effects are necessary in order to conclusively demonstrate a functional distinction among regions (Nieuwenhuis et al., 2011). Such interactions have rarely been shown, limiting the ability to claim strong differences in function among regions (for discussion, see Blank et al., 2016; Blank & Fedorenko, 2020; Fedorenko et al., 2020; cf. Matchin & Wood, 2020). Moreover, many neuroimaging experiments have actually shown similar activations in language-related cortex for different aspects of linguistic processing, particularly for syntax and semantics (Diachek et al., 2020; Fedorenko, Nieto-Castañon, et al., 2012; Fedorenko et al., 2020; Humphries et al., 2006, 2007; Matchin et al., 2017, 2019).

In response to this lack of conclusive evidence, some researchers have questioned whether there is in fact functional segregation across regions of the language network involved in lexical access, syntactic processing, and semantic interpretation, instead advocating for a shared processing mechanism (I. Blank et al., 2016; I. A. Blank & Fedorenko, 2020; Fedorenko et al., 2020; Mahowald & Fedorenko, 2016). This idea takes inspiration from linguistic theory, which posits a close interconnection among these systems. However, the fact that regions of the language network often show a somewhat similar activation profile in neuroimaging experiments does not distinguish between a neural architecture with the same functional mechanism across regions and a neural architecture with tightly interconnected yet distinct mechanisms across regions. This is because lexical, syntactic, and semantic components of language are systematically connected. For example, inverting the order of words in a sentence like *dog bites man* results in a far more surprising semantic interpretation than the original. Thus, any experimental manipulation of one component is likely to affect another component, resulting in similar neuroimaging effects across regions, despite the possibility that these regions in fact underlie distinct mechanisms.

This tight connection between components of language poses an obstacle to identifying the potentially distinct brain bases of higher-level linguistic functions through the use of neuroimaging. *Lesion-symptom mapping* (LSM), the study of associations between brain damage and behavioral deficits, can help resolve this conundrum. Lesion-symptom mapping allows a researcher to identify brain regions necessarily involved in a particular linguistic function, rather than functional neuroimaging in healthy individuals, which only provides correlative information (Bates et al., 2003; Rorden & Karnath, 2004; Wilson, 2017). Previous lesion-symptom mapping studies have associated different language-related brain regions with different linguistic processes (J. Ding et al., 2020; Dronkers et al., 2004; Gleichgerrcht et al., 2016; Kristinsson et al., 2020; Magnusdottir et al., 2013; Matchin et al., 2020; Mesulam et al., 2015; Pillay et al., 2017; Riccardi et al., 2020; Rogalsky et al., 2018; Schwartz et al., 2011; Thothathiri et al., 2012; Wilson, Henry, et al., 2010; Wilson, Dronkers, et al., 2010). However, it could be the case that anatomical or other forms of variability could result in a significant lesion-deficit association in one area and a subthreshold association in another, but this does not mean that the first region is significantly *more* implicated in such deficits than the second. As with functional neuroimaging, region by condition interaction effects are necessary to show that some region is more strongly implicated in a given task than another region, but none of these previous studies report task by region interaction effects. Therefore, while it is plausible that different regions process different aspects of language, it has not yet been conclusively shown using region by condition interaction analyses, which directly compare the strength of effects between regions.

In order to address this issue, we performed a LSM study assessing measures tapping into distinct linguistic processes within the broader categories of syntactic and conceptual-semantic processing. We focused on syntax and semantics because functional neuroimaging studies aiming to identify distinct neurobiological bases for these domains in sentence comprehension have frequently found very tight overlap of syntactic and semantic effects among all frontal-temporal-parietal regions implicated in language (Fedorenko, Nieto-Castañon, et al., 2012; Fedorenko et al., 2020; Matchin et al., 2017). We do not believe there has yet been offered a perfect decomposition of the set of all syntactic and conceptual-semantic mechanisms involved in language. However, we were guided in our analyses by the theoretical model that two of us recently published that ascribes distinct syntactic and conceptual-semantic functions to different regions of the language network (Matchin & Hickok, 2020). This model posits that conceptual-semantic processing, equally for both comprehension and production, is primarily supported by two regions: the anterior superior temporal sulcus (aSTS) and inferior angular gyrus (iAG). The model posits that syntactic processing is primarily supported by two different regions, differentially for comprehension and production: the posterior middle temporal gyrus (pMTG) supports hierarchical syntactic structure building necessary for comprehension and production, and the posterior inferior frontal gyrus (pIFG) supports morpho-syntactic processes necessary for production but not for comprehension.

We performed our analyses based on a large extant database of subjects and tasks that were not designed to perfectly isolate the syntactic and semantic mechanisms identified in the Matchin & Hickok model. However, we predicted that our selected measures would allow us to identify significant measure by region interaction effects. Given the predictions of the model, we posited that we would find that:

- damage to pMTG would be more significantly associated with syntactic comprehension deficits than pIFG;
- damage to pIFG would be more significantly associated with deficits in morpho-syntactic production, that is expressive agrammatism, than aSTS;
- damage to iAG would be more significantly associated with deficits in conceptual-semantic retrieval than pIFG, regardless of whether this is assessed in production or comprehension.

Finding such interaction effects would support the Matchin & Hickok model as well as theories of language organization in the brain that posit distinct syntactic and conceptual-semantic functions in different regions of the language network more generally, and would cast doubt on recent hypotheses that syntax and semantics are processed jointly in a unified function across all frontal-temporal-parietal regions of the language network.

## 2 Materials & Methods

### 2.1 Subjects & Measures

In three partially overlapping groups of subjects, we assessed four different measures: Group 1, N = 121, *Syntactic Comprehension*; Group 2, N = 92, *Expressive Agrammatism* and *Semantic Category Word Fluency*; and Group 3, N=218, *Word Comprehension*. Subjects were assessed on a number of language batteries, which were part of multiple studies on aphasia recovery. Group 1 subjects were the same as reported in Den Ouden et al. (2019) and Kristinsson et al. (2020). 47 subjects were included solely in Den Ouden et al. (2019), 48 subjects were included solely in Kristinsson et al. (2020), and 26 subjects were included in both studies. Group 2 subjects were the same as reported in Den Ouden et al., (2019) and Matchin et al., (2020). 39 subjects were included solely in Den Ouden et al. (2019), 32 were included solely in Matchin et al. (2020), and 21 subjects were included in both studies (for the 21 subjects that were included in both studies, we used the ratings in Matchin et al., 2020). All of the lesion maps and behavioral data for subjects enrolled in this study are available for download at https://www.dropbox.com/sh/3w4aeizgypfs7sd/AAB-W8Yn5qDUFeBj90WKsBqAa?dl=0.

All subjects were recruited through local advertisement. They provided informed consent to participate in this study, which was approved by the Institutional Review Boards at the University of South Carolina and the Medical University of South Carolina. All subjects had at least one ischemic stroke to the left hemisphere at least six months prior to study inclusion and were also pre-morbidly right-handed (self-disclosed). Demographic information for the three groups of subjects is shown in Table 1.

**Table 1.** Subject information for the two partially overlapping groups of subjects. SD = standard deviation. AQ = aphasia quotient of the Western Aphasia Batter-Revised, a summary measure of overall language ability. * education information was available for only 210/218 of these subjects.

|  | **Group 1** | **Group 2** | **Group 3** |
| --- | --- | --- | --- |
| *Measures* | Syntactic Comprehension | Semantic Category Word Fluency<br>Expressive Agrammatism | Word Comprehension |
| *Total number of subjects* | N = 121 | N = 92 | N = 218 |
| *Sex* | 79 male, 42 female | 60 male, 32 female | 133 male, 85 female |
| *Mean age at initial testing (years)* | 59.6 (SD = 10.6) | 58.7 (SD = 11.1) | 60.0 (SD = 11.4) |
| *Mean months post-first stroke at initial testing* | 47.5 (SD = 52.4) | 46.4 (SD = 49.6) | 43 (SD = 48.4) |
| *Mean number of strokes* | 1.17 (SD = 0.49) | 1.11 (SD = 0.43) | 1.17 (SD = 0.48) |
| *Mean education (years)* | 15.4 (SD = 2.3) | 15.3 (SD = 2.3) | 15.0 (SD = 2.3)* |
| *Mean lesion volume (mm<sup>3</sup>)* | 112,652 (SD = 94,945) | 96,175 (SD = 84,514) | 120,855 (SD = 97,488) |
| *Mean WAB-R AQ* | 66.0 (SD = 26.9) | 75.8 (SD = 19.9) | 61.4 (SD = 28.1) |

The *Syntactic Comprehension* measure was designed to assess the syntactic processes that are necessary for assigning a hierarchical syntactic structure to an incoming sentence in order to correctly interpret its meaning. We derived this measure from two different sentence-picture matching tasks reported in more detail elsewhere (Den Ouden et al., 2019; Kristinsson et al., 2020), with lesion data for subjects included here partially reported in Den Ouden et al. (2019) and Fridriksson et al. (2018). The tasks involved a range of constructions, but our focus here is on complex, semantically reversible sentences with non-canonical word order. These included object-extracted clefts (Kristinsson et al., 2020; e.g., *it is the boy that the girl chases*), object-extracted relative clauses (Den Ouden et al., 2019; e.g., *the boy that the girl chased is happy*), and object-extracted Wh-questions (both studies; e.g., *which boy did the girl chase?*). Sentences of this sort have a long history in research on syntactic ability in comprehension because (i) lacking canonical English subject-verb-object word order and (ii) lacking semantic plausibility constraints (c.f., *which apple did the boy eat?*), they require syntactic analysis for determining who is doing what to whom (Caramazza & Zurif, 1976). Performance on such sentences is standardly compared to sentences with canonical word order containing the same verbs and nous (i.e., the noun verbed the noun) as a control for speech perception, lexical processing, basic ability to infer an event structured based on a sequence of words, processing of semantic relations, and working memory or decision-making resources involved in performing a sentence-picture matching task (Caramazza & Zurif, 1976; Cho-Reyes & Thompson, 2012; Love & Oster, 2002; Rogalsky et al., 2018; Thompson et al., 2013; Thothathiri et al., 2012). Accordingly, we created our syntactic comprehension measure as the average performance on complex, noncanonical sentences with performance on simple, semantically reversible active sentences covaried out using linear regression. Importantly, the same verbs and agent and patient nouns (and thus the same thematic relations) were included in both the active and non-canonical sentence types to control for lexical and relational semantics. For subjects who performed both studies (N = 26), in order to provide the most reliable estimate, scores were averaged across the two studies, which contained the same number of trials (5 trials per sentence type).

The *Expressive Agrammatism* measure was designed to assess the morphosyntactic processes that are necessary for overtly expressing a message, which two of us have recently argued to be separable from the hierarchical syntactic processing that is necessary for sentence comprehension (Matchin & Hickok, 2020). We derived this measure from samples of connected speech production elicited either by (i) describing the Cookie Theft picture (Goodglass & Kaplan, 1983, as reported in Den Ouden et al., 2019) or (ii) retelling the story of Cinderella in their own words (MacWhinney et al., 2011, as reported in Matchin et al., 2020). The presence of expressive agrammatism was determined as described in Matchin et al., (2020). Briefly, production samples were rated independently by speech and language experts for the systematic simplification of sentence structure and omission of function words and morphemes. This resulted in a categorical assessment for each subject, either agrammatic or not. Given that categorical, binary ratings of agrammatism were used in Den Ouden et al. (2019) and Matchin et al. (2020), we did not average scores across these two studies. Given that agrammatic patients tend to have slower, more effortful speech (Damasio, 1992; Goodglass & Kaplan, 1983), we included speech rate as a covariate using logistic regression (words per minute during the task) in order to focus on residual morphosyntactic production abilities rather than general articulatory fluency.

The *Semantic Category Word Fluency* measure was designed to assess the retrieval of conceptual-semantic content associated with words, which is separable from the syntactic processes described above which are associated with the form, rather than the meaning, of sentences. This measure came from the Word Fluency subtest of the Western Aphasia Battery - Revised (WAB-R) (Kertesz, 2007), as administered by a licensed speech language pathologist. Because subjects were given a highly variable number of WAB-R assessments, we selected the first available WAB-R for each subject. The Word Fluency subtest involves asking the subject to name as many animals as possible within one minute (maximum score is 20). Word fluency tasks for semantic categories are generally is designed to assess two broad categories of abilities: access to conceptual-semantic representations and executive function (Chertkow & Bub, 1990; Troyer et al., 1997; Unsworth et al., 2011). However, Whiteside et al. (2016) performed a factor analysis of a highly similar semantic category word fluency task and found that deficits on this task were associated with language measures but not executive function measures. Furthermore, lesion-symptom mapping (Baldo et al., 2006) and functional neuroimaging (Birn et al., 2010) studies of semantic category word fluency implicate the inferior angular gyrus, a region strongly associated with conceptual-semantic processing, broadly construed (Binder et al., 2009; Hodgson et al., 2021; Humphreys et al., 2021; Lau et al., 2008), among other regions also associated with semantic processing. The lesion-symptom mapping study of Baldo et al. found that only a letter fluency task was associated with frontal lesions, not semantic category word fluency. These results suggest that semantic word fluency tasks, at least for tasks with broad semantic categories like animals, load highly on semantic processing and less highly on executive function, consistent with the fact that patients with semantic dementia perform worse on semantic category fluency measures than letter fluency relative to patients with Alzheimer’s disease (Marczinski & Kertesz, 2006). To control for articulatory fluency, we incorporated the same speech rate covariate we used for the expressive agrammatism analysis.

The *Word Comprehension* measure was designed to provide an alternative window into conceptual-semantic retrieval processes. Most models of language in the brain, including the Matchin & Hickok model, postulates that regions involved in conceptual-semantic processing support both comprehension and production equally. Because the *Semantic Category Word Fluency* measure involves speech production, we chose the *Word Comprehension* measure to assess conceptual-semantic retrieval in speech comprehension. The measure came from the auditory word comprehension subtest of the Western Aphasia Battery - Revised (WAB-R) (Kertesz, 2007), as administered by a licensed speech language pathologist. The Auditory Word Recognition subtest involves verbally requesting the subject to point to printed images or real-world objects. The experimenter prompts subjects with a sentence, e.g., “point to the __” or “show me the __”. The test involves multiple types of tested words, including real household objects (cup, matches, pencil, flower, comb, screwdriver), pictured objects (the same as real objects), pictured shapes (square, triangle, circle, arrow, cross, cylinder), pictured letters (J, F, B, K, M, D), pictured numbers (5, 61, 500, 1867, 32, 5000), pictured colors (blue, brown, red, green, yellow, black), real world furniture (window, chair, desk or bed, light, door, ceiling), real world body parts (ear, nose, eye, chest, neck, chin), real world fingers (thumb, ring finger, index finger, little finger, middle finger), and real world body parts on the correct side (right ear, right shoulder, left knee, left ankle, right wrist, left elbow, right cheek). For each item the subject receives 1 point, for a total of 60 points.

### 2.2 Brain Imaging & Lesion mapping

We acquired anatomical MRIs and performed lesion mapping using the same parameters and procedures as described in Fridriksson et al. (2018). Neuroimaging data were collected at the University of South Carolina and the Medical University of South Carolina. Lesions were demarcated onto each subject’s T2 image by an expert technician or an expert neurologist blind to the behavioral data.

Lesion overlap maps for both groups are shown in Figure 1. Overall, there was good coverage in perisylvian cortex, covering all selected regions of interest (described below).

**Figure 1.**
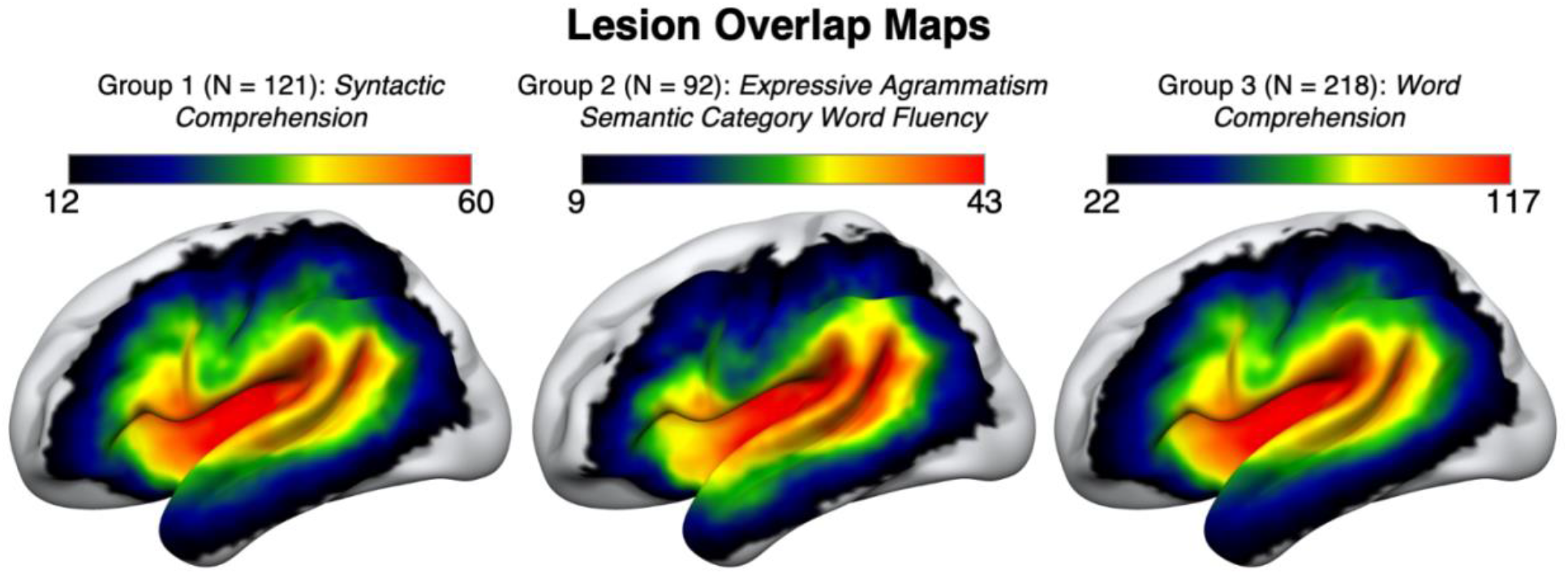
Lesion overlap maps for each of the groups. LEFT: Group 1 (N = 121), assessed for *Syntactic Comprehension*. The lower boundary of 12 corresponds to voxels where at least 10% of subjects had damage. MIDDLE: Group 2 (N = 92), assessed for *Semantic Category Word Fluency* and *Expressive Agrammatism*. The lower boundary of 9 corresponds to voxels where at least 10% of subjects had damage. RIGHT: Group 3 (N = 218), assessed for *Word Comprehension*. The lower boundary of 22 corresponds to voxels where at least 10% of subjects had damage.

### 2.3 Region of Interest (ROI) definition

Rather than using structurally-defined ROIs, which may not line up precisely with linguistically-relevant regions of the brain of interest in the present study, we used the statistical maps associated with a previous fMRI study on sentence processing (Matchin et al., 2017) to define ROIs for analysis. This study compared multiple conditions, including full natural sentences (e.g., *the poet might recite a verse*) and jabberwocky sentences, which involve the substitution of pseudowords for content words (e.g., *the tevill will sawl a pand*). The contrast of natural sentences > jabberwocky sentences highlighted a number of language-related brain regions in association cortex that are frequently identified in brain imaging studies of syntax and semantics. Importantly, this contrast ensured adequate coverage of all regions of interest, whereas similar contrasts of structure (e.g., sentences compared to word lists) produced very little extent of activation in iAG and relatively smaller extent of activation the posterior temporal lobe. We therefore used the natural sentences > jabberwocky sentences contrast with a reduced voxel-wise threshold of p < 0.01 to ensure adequate ROI size and coverage for brain regions that have been implicated in both syntactic and semantic processing (Figure 2, LEFT). We selected clusters corresponding to four left hemisphere regions that have been previously implicated in these processes (Figure 2, RIGHT): the inferior angular gyrus (iAG), the posterior middle temporal gyrus (pMTG), the anterior superior temporal sulcus (aSTS), and the posterior inferior frontal gyrus (pIFG). Several of the clusters revealed by the analysis were contiguous at local minima, and so we manually separated them at these junctures to form four separate ROIs.

**Figure 2.**
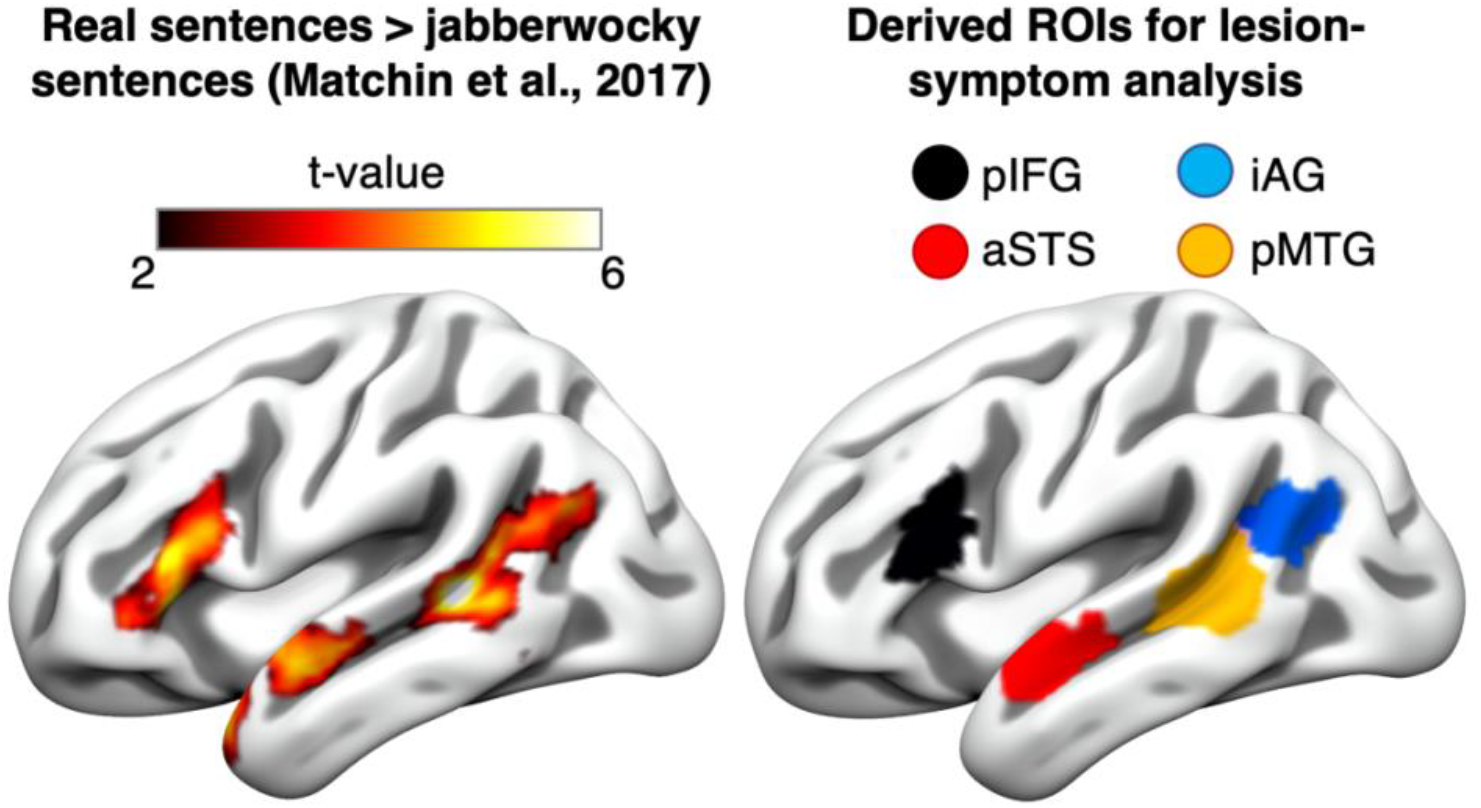
Regions of interest (ROIs) derived from previously published functional neuroimaging data. LEFT: Statistical contrast of natural sentences > jabberwocky sentences from Matchin et al., (2017) at the uncorrected voxel-wise threshold of p < 0.01. RIGHT: Selected ROIs using the clusters derived from the contrast of natural sentences > jabberwocky sentences from Matchin et al., (2017). Blue: inferior angular gyrus (iAG); orange: posterior middle temporal gyrus (pMTG); red: anterior superior temporal sulcus (aSTS); black: posterior inferior frontal gyrus (pIFG).

### 2.4 Lesion Analyses

In order to assess the overall relationship between our behavioral measures and damage to language-relevant regions, we performed ROI-based univariate and multivariate regression analyses in NiiStat (https://www.nitrc.org/projects/niistat/) using the set of four ROIs we derived from Matchin et al. (2017). Some authors have pointed out the spatial distortion in univariate lesion-symptom mapping analyses that occurs based on the non-random distribution of lesions in the brain and the potential superiority of multivariate methods in reducing this distortions (Mah et al., 2014). However, Ivanova et al. (2021) pointed out that these comparisons involved outdated and procedures for univariate analyses. They performed a systematic comparison of univariate and multivariate approaches to lesion-symptom mapping, incorporating crucial features that were absent from Mah et al. (2014): permutation testing for multiple comparisons, lesion volume as a covariate, and minimum 5-10% lesion load threshold. Ivanova et al. (2021) found that, with these updated procedures, univariate methods actually outperformed multivariate methods on most dimensions related to spatial distortion. They recommend reporting both univariate and multivariate analyses incorporating these crucial procedures. We accordingly corrected for multiple comparisons using permutation tests (10,000 permutations), with a minimum lesion load of 10% of sample, and all analyses incorporated lesion volume as a covariate, as also recommended by DeMarco & Turkeltaub (2018). We supplement these results with unthresholded univariate voxel-wise lesion maps associated with each measure in order to illustrate that our ROI analyses did not obscure the lesion distribution associated with each measure, in a similar fashion as is recommended for fMRI analyses (Poldrack et al., 2008).

We also performed four hypothesis-driven interaction analyses, one for each measure. With respect to *Syntactic Comprehension*, we tested the interaction between the pMTG and pIFG. Both the pIFG and pMTG are commonly activated in neuroimaging studies of syntactic comprehension. While most theoretical models of syntax posit a key role for the pIFG in receptive syntax (Friederici, 2017; Hagoort, 2014; Tyler & Marslen-Wilson, 2008), previous lesion-symptom mapping studies have indicated that damage to posterior temporal areas (but not frontal areas) is critically implicated in syntactic comprehension deficits (Kristinsson et al., 2020; Pillay et al., 2017; Rogalsky et al., 2018). In this light, the models proposed by Matchin & Hickok (2020) and Bornkessel-Schlesewsky & Schlesewsky (2013) posit that the pIFG is not critically involved in syntactic comprehension. Therefore, we hypothesized that damage to the pMTG would be significantly more implicated in syntactic comprehension deficits than damage to the pIFG.

Most theoretical models of syntax in the brain attribute a key role in syntactic production to the pIFG but not the aSTS (Friederici, 2017; Hagoort, 2014; Matchin & Hickok, 2020; Tyler & Marslen-Wilson, 2008). Consistent with this, agrammatism is primarily associated with damage to inferior frontal cortex, and to a lesser extent posterior temporal-parietal cortex, but not anterior temporal cortex (Sapolsky et al., 2010; Wilson et al., 2010; Den Ouden et al., 2019; Matchin et al., 2020). Therefore, with respect to our *Expressive Agrammatism* measure, we tested the interaction between aSTS and pIFG. We expected that our measure would be significantly more associated with damage to pIFG than aSTS.

Finally, with respect to *Semantic Category Word Fluency*, we tested the interaction between iAG and pIFG. Although damage to both of these regions has been claimed to be implicated in lexical-semantic deficits (Fedorenko et al., 2020), damage to iAG and surrounding temporal cortex, but not frontal cortex, was previously shown to be associated with deficits on a similar word fluency measure (Baldo et al., 2006). Consistent with this, most theories attribute a (morpho-)syntactic function to the pIFG (Friederici, 2017; Hagoort, 2014; Matchin & Hickok, 2020; Tyler & Marslen-Wilson, 2008), or a top-down selection mechanism (Novick et al., 2005; Thompson-Schill & Cutler, 2005), but not a basic lexical or conceptual-semantic function. Therefore, we expected that damage to iAG would be significantly more associated with impairments on this measure than damage to pIFG.

To bolster our test of the interaction between iAG and pIFG with respect to conceptual-semantic processing, we tested the same interaction using the *Word Comprehension* measure. Unlike *Semantic Category Word Fluency*, which involves speech production and likely includes some degree of an executive function component, our word comprehension measure does not require speech output, and minimizes executive function demands. However, like *Semantic Category Word Fluency*, it involves accessing an item in the lexicon and its associated conceptual-semantic information. Previous studies on word comprehension, without including a covariate for non-linguistic conceptual knowledge, have found an association between impaired performance and damage primarily to temporal and inferior parietal lobe regions (Fridriksson et al., 2018; Hart & Gordon, 1990; Hillis et al., 2001; Selnes et al., 1983). Therefore, we expected that deficits on this task would be associated with damage to iAG like for *Semantic Category Word Fluency* (in addition to pMTG and possible aSTS). We similarly expected an association between *Word Recognition* deficits and damage to iAG relative to pIFG as we expected for *Semantic Category Word Fluency*.

To test these interactions, we first calculated proportion damage to each ROI and adjusted the data using a rationalized arcsine transform (Studebaker, 1985), and then computed residual damage values by covarying out the effect of lesion volume. We then assessed the region by measure interaction effect in linear regression for each of the three measures of interest in SPSS. We corrected for multiple comparisons using a Bonferroni correction with an adjusted alpha threshold of p < 0.0125 for each of the four comparisons, controlling the total error at p < 0.05.

## 3 Results

The univariate ROI analyses, corrected for multiple comparisons (FWE p < 0.05), revealed the following effects:

- *Syntactic comprehension*: two ROIs showed a significant negative association between percent damage and behavioral scores, pMTG (Z = −3.13) and aSTS (Z = −2.46).
- *Expressive agrammatism*: one ROI showed a significant positive association between percent damage and behavioral scores, pIFG (Z = 3.10). I.e., subjects who had stronger expressive agrammatism scores were more likely to have damage to pIFG.
- *Word Comprehension:* three ROIs showed a significant negative association between percent damage and behavioral scores, iAG (Z = −3.50), pMTG (Z = −3.39), and aSTS (−2.18).
- *Semantic category word fluency*: one ROI showed a significant negative association between percent damage and behavioral scores, iAG (Z = −3.91).

The multivariate ROI analyses, corrected for multiple comparisons (FWE p < 0.05), revealed the following effects:

- *Syntactic Comprehension*: two ROIs showed a significant negative association between percent damage and behavioral scores, pMTG (Z = −2.07) and aSTS (Z = −1.95).
- *Expressive Agrammatism*: no ROI showed a significant positive association between percent damage and behavioral scores.
- *Word Comprehension*: two ROIs showed a significant negative association between percent damage and behavioral scores, iAG (Z = −2.46) and pMTG (Z = −2.35).
- *Semantic Category Word Fluency*: one ROI showed a significant negative association between percent damage and behavioral scores, iAG (Z = −2.20).

Unthresholded univariate voxel-wise maps support the localization of the effects revealed by the ROI analyses, (Figure 3), although the strongest effect of *expressive agrammatism* was in the posterior inferior frontal sulcus/middle frontal gyrus rather than the IFG itself.

**Figure 3.**
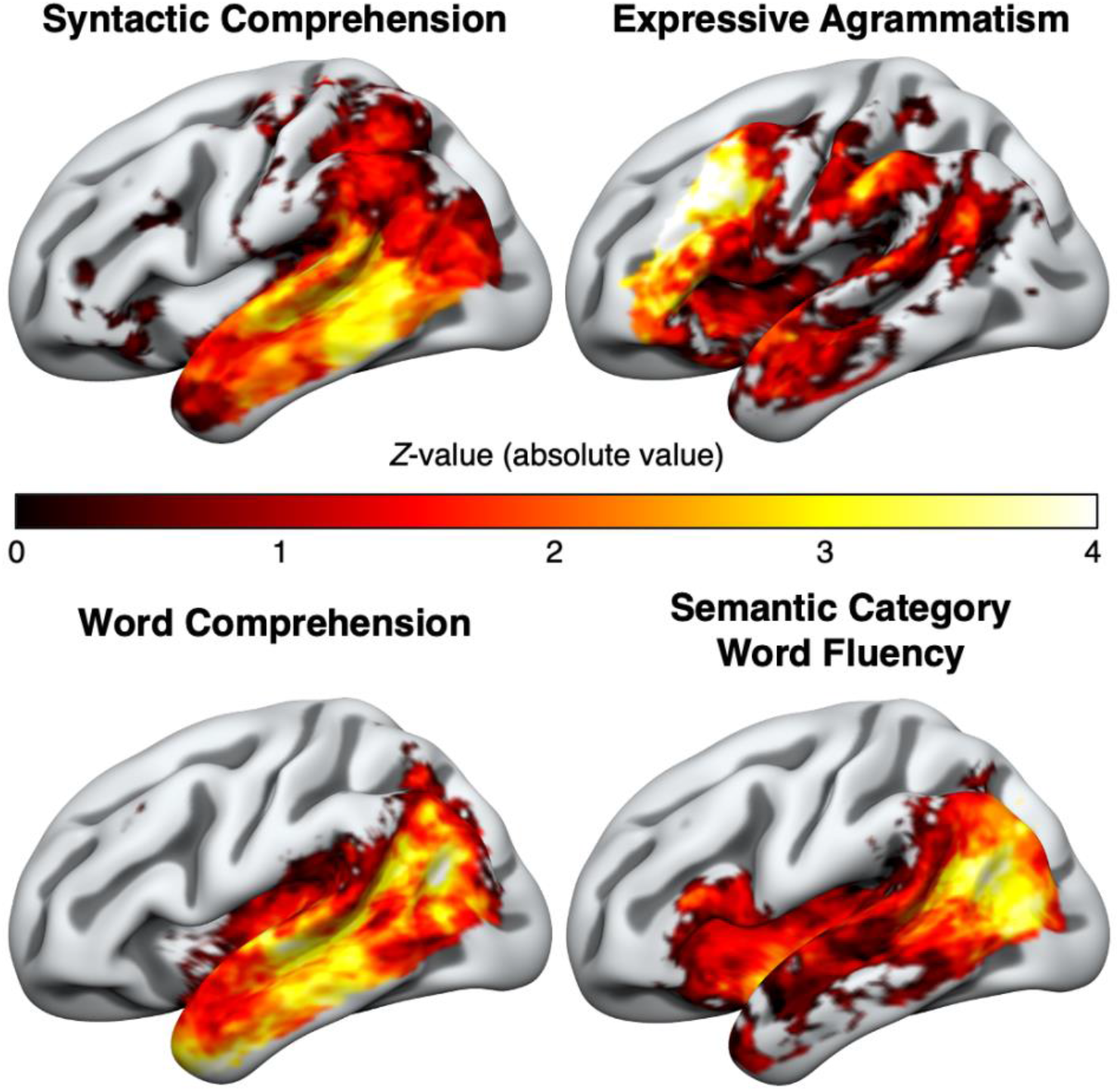
Unthresholded voxel-wise univariate analyses (shown in absolute value of Z-scores) for each of the four behavioral measures. UPPER LEFT: *Syntactic Comprehension* (noncanonical sentence comprehension performance with active sentence comprehension as a covariate), UPPER RIGHT: *Expressive Agrammatism* (perceptual ratings of agrammatism with speech rate as a covariate), BOTTOM LEFT: *Word Comprehension* (), and *Semantic Category Word Fluency* (WAB word fluency with speech rate as a covariate). Each measure is identified at top in bold, corresponding to all of the figures underneath.

Scatterplots illustrating the *region by measure* interactions we tested are shown in Figure 4. Cohen (1988) recommends interpreting effect sizes (*η*^2^) with the following benchmarks: 0.01=small; 0.06=medium; 0.14=large.

**Figure 4.**
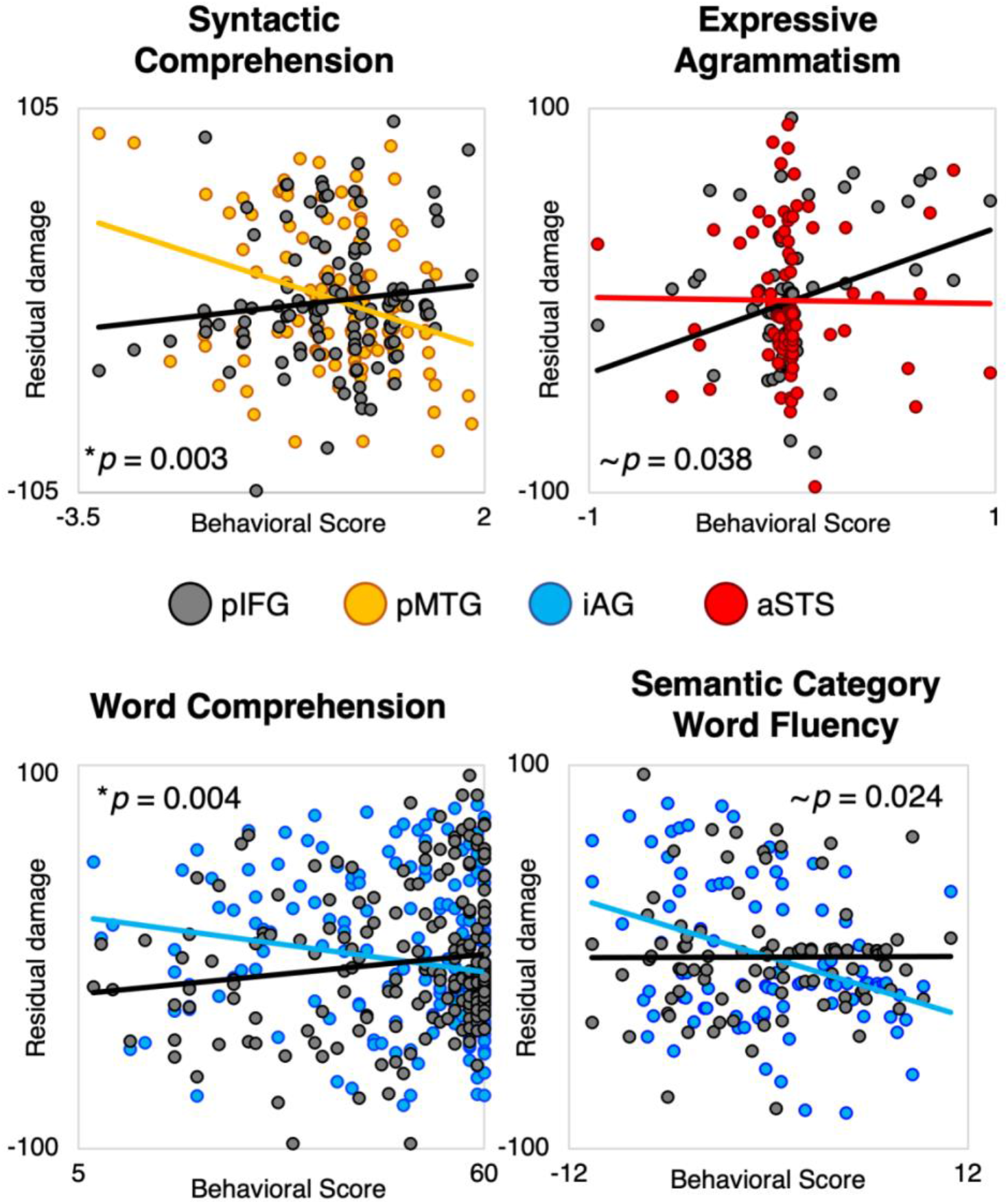
Scatter plots illustrating four hypothesis-driven *region by measure* interaction analyses, corrected for multiple comparisons using a Bonferroni correction (individual p < 0.0125), for the four ROIs of interest (pIFG, pMTG, iAG, aSTS) and the four behavioral measures. TOP LEFT: *Syntactic Comprehension* (noncanonical sentence comprehension performance with active sentence comprehension as a covariate), TOP RIGHT: *Expressive Agrammatism* (perceptual agrammatism ratings with speech rate as a covariate), BOTTOM LEFT: Auditory Word Comprehension (WAB auditory word comprehension subscore), and BOTTOM RIGHT: (*Semantic Category Word Fluency* (WAB word fluency with speech rate as a covariate).

- *Syntactic Comprehension*: the association between deficits on this measure with damage to the pMTG was significantly stronger than with damage to the pIFG, F(1,119) = 9.013, *p* = 0.003, *η*^2^ = 0.076.
- *Expressive Agrammatism*: the association between positive assessment on this measure with damage to the pIFG was marginally significantly stronger than with damage to the aSTS, F(1,90) = 4.432, *p* = 0.038, *η*^2^ = 0.023. I.e., subjects who had stronger expressive agrammatism scores were more likely to have damage to pIFG than to aSTS.
- *Auditory Word Comprehension:* the association between deficits on this measure with damage to the iAG was significantly stronger than with damage to the pIFG, F(1,216) = 8.440, *p* = 0.004, *η*^2^ = 0.039.
- *Semantic Category Word Fluency*: the association between deficits on this measure with damage to the iAG was marginally significantly stronger than with damage to the pIFG, F(1,90) = 5.294, *p* = 0.024, *η*^2^ = 0.040.

Residual damage indicates the residual percent damage values within an ROI, with lesion volume as a covariate. Straight lines indicate estimated linear trends. pMTG = posterior middle temporal gyrus; pIFG = posterior inferior frontal gyrus; iAG = inferior angular gyrus; aSTS = anterior superior temporal sulcus. * significant effect. ∼ effect approaching significance.

## 4 Discussion

In this lesion-symptom mapping (LSM) study in three groups of patients with chronic post-stroke aphasia (N=218, N=121, N=92), deficits in four distinct measures of linguistic processing were each associated with distinct patterns of damage within language network: syntactic comprehension was associated primarily with pMTG damage and secondarily with aSTS damage, expressive agrammatism was associated with pIFG damage, word comprehension was associated with iAG, pMTG, and aSTS damage, and semantic category word fluency was associated with iAG damage. None of these effects are unique to our study, supporting similar previous findings in the literature, as we discuss below. However, critically, we also showed region by measure interaction effects, such that damage to specific regions in the language network was more associated with each behavioral measure than damage to other regions, significantly or trending toward significance.

Combined, these results narrow down possible functions of these brain regions in higher-level linguistic processing, and suggest that neuroimaging research needs to incorporate insights from lesion symptom mapping in order to understand the architecture of language in the brain. Namely, the region by measure interactions we identified present strong challenges to the hypothesis that language-related brain regions underlie a highly similar or identical function with respect to linguistic processing, a hypothesis that has been advocated most recently in light of neuroimaging data alone (I. Blank et al., 2016; I. A. Blank & Fedorenko, 2020; Caplan et al., 1996; Dick et al., 2001; Fedorenko et al., 2020; Mollica et al., 2020). Our results are instead broadly consistent with neuroanatomical models that posit distinct linguistic functions to different regions within the language network (Bornkessel-Schlesewsky & Schlesewsky, 2013; Friederici, 2017; Hagoort, 2014; Matchin & Hickok, 2020; Tyler & Marslen-Wilson, 2008). However, the model presented by Matchin & Hickok (2020) provides explanations for all of the identified effects and interactions, whereas the other models fail to provide clear explanations for one or more of them. We discuss each of these measures in turn with respect to existing lesion-symptom mapping literature and how these findings relate to existing models of language organization in the brain.

### 4.1 Expressive agrammatism

The association between the classical, production-related deficit of agrammatism and damage to inferior frontal cortex has a long history stemming back to the origins of aphasiology (Kleist, 1914; Kussmaul, 1877; Pick, 1913; Tissot et al., 1973), supported by recent LSM studies (Sapolsky et al., 2010; Wilson et al., 2010; Den Ouden et al., 2019; Matchin et al., 2020). Our results, combining data from two previously reported studies (Den Ouden et al., 2019; Matchin et al., 2020), reaffirm this association. The anterior temporal lobe (ATL) more broadly has been associated with syntax (i.e., comprehension of sentences or phrases vs. word lists) in many neuroimaging studies (Bemis & Pylkkanen, 2011; J. Brennan et al., 2012; J. R. Brennan et al., 2016; Humphries et al., 2005, 2006; Mazoyer et al., 1993; Rogalsky et al., 2011; Rogalsky & Hickok, 2009). However, previous research on patients with ATL damage and/or degeneration shows no evidence of agrammatic production deficits (Hodges et al., 1992; Hodges & Patterson, 2007; Kho et al., 2008; Mesulam et al., 2015; Corianne Rogalsky et al., 2018; see Wilson et al., 2014 for data and a review). Our study supports the general picture painted by previous lesion-symptom mapping studies in showing that the association of expressive agrammatism with pIFG damage was stronger than with aSTS damage, trending toward significance. This suggests that the apparent involvement of the ATL in syntax indicated by some neuroimaging studies may in fact be due to semantic processing downstream from syntax, rather than syntax per se (Pallier et al., 2011; Wilson et al., 2014).

This suggests the existence of some syntactic mechanism involved in sentence production associated specifically with the pIFG but not the aSTS. While most theories of language organization in the brain posit a syntactic mechanism in inferior frontal cortex (Friederici, 2017; Hagoort, 2014; Matchin & Hickok, 2020; Tyler & Marslen-Wilson, 2008), the theory proposed by Bornkessel-Schlesewsky & Schlesewsky (2013) does not, suggesting that the pIFG supports linguistic processing through a more general cognitive control function. However, this theory attributes syntactic processing to a dorsal stream spanning posterior temporal cortex through parietal and frontal cortex. On this point, the fact that our analysis only approached significance (p = 0.038, corrected threshold p < 0.0125) may be due to the fact that the posterior middle frontal cortex and not pIFG was most strongly associated with expressive agrammatism. Future research should further investigate the role of middle frontal regions in linguistic processing.

### 4.2 Syntactic comprehension

We found that damage to the pMTG was significantly more associated with syntactic comprehension deficits compared to the pIFG, which was not implicated at all. Previous LSM studies have found an association between residual syntactic comprehension scores (regressing out single word comprehension or production) with damage to the posterior temporal lobe but not the pIFG (Kristinsson et al., 2020; Pillay et al., 2017; Rogalsky et al., 2018). Our results reinforce these previous studies by showing that the effect in pMTG is statistically stronger than in pIFG (in fact, the pIFG showed an effect numerically in the opposite direction). As with expressive agrammatism, this result is incompatible with the shared-mechanism view of the language network. Most models of language organization in the brain posit that the pIFG is not only critical for processing hierarchical structure in production but also comprehension (Friederici, 2017; Hagoort, 2014; Tyler & Marslen-Wilson, 2008). However, the data are more consistent with the proposals of Matchin & Hickok (2020) and Bornkessel-Schlesewsky & Schlesewsky (2013), who argue instead that the pIFG’s role in sentence comprehension is restricted to a supporting mechanism, but not critical for building hierarchical structure.

Our univariate analyses also identified a significant, though weaker, association between syntactic comprehension deficits and damage to the aSTS. Some LSM studies have found some evidence of an association between syntactic comprehension deficits and ATL damage (Dronkers et al., 2004; Magnusdottir et al., 2013). Matchin et al. (2020) speculated that these effects might have been due to the lack of lesion volume as a control, predicting that including lesion volume might eliminate such an association. However, the analyses reported here all included lesion volume as a covariate, suggesting that syntactic comprehension deficits may be associated with anterior temporal as well as posterior temporal damage. This is somewhat similar to previous studies of syntactic comprehension deficits, which generally found more robust effects in posterior temporal cortex but some association with noncanonical sentence comprehension deficits in aSTS (Pillay et al., 2017; Rogalsky et al., 2018; Kristinsson et al., 2020). This raises the question of whether the aSTS is involved in some aspect of combinatorial processing, which may be semantic in nature (Pylkkänen, 2020). Future studies should seek to investigate possible function segregation between pMTG and aSTS along these lines. We hypothesize that semantic combination (controlling for syntax) will be more associated with aSTS damage, whereas syntactic combination (controlling for semantics) will be more associated with pMTG damage.

### 4.3 Semantic category word fluency

In our univariate analyses, damage to the iAG and no other region was significantly associated with deficits on the semantic category word fluency measure. Furthermore, our region by measure interaction analysis revealed that damage to the iAG was more associated with deficits than damage to the pIFG, trending towards significance. Our result is similar to that of Baldo et al. (2006), who found that deficits on a semantic similar word fluency task were associated with posterior temporal and inferior parietal damage but not frontal damage.

Most theories of language organization in the brain posit a role for lexical and/or conceptual-semantic processing in the iAG (Friederici, 2017; Hagoort, 2014; Tyler & Marslen-Wilson, 2008; Matchin & Hickok, 2020; cf. Bornkessel-Schlesewsky & Schlesewsky, 2013). Many neuroimaging studies have indicated that this region is particularly responsive to manipulations of semantics, but not syntax per se independently of conceptual content (see Matchin & Hickok, 2020 for a review). For example, Pallier et al. (2011) showed that this region responded to increased linguistic structural complexity, but not when meaningful content words were replaced with pseudowords, unlike other regions (namely, the pMTG and anterior IFG) that responded to structural complexity regardless of meaningfulness. This dissociation has been replicated multiple times (Fedorenko, Nieto-Castañon, et al., 2012; Goucha & Friederici, 2015; Matchin et al., 2017). This suggests that rather than processing the hierarchical syntactic structure of a sentence itself, this region processes the complex semantic representation that results from syntactic combination. Similarly, Price et al. (2015) found that processing meaningful word pair combinations (e.g. “plaid jacket”) results in more activity in iAG than less meaningful combinations (e.g. “moss pony”), including controls for co-occurrence frequency. Our results, an association between iAG damage and deficits in semantic category word fluency, are strongly consistent with this literature.

According to the view of undifferentiated higher-level linguistic processing, pIFG should have also been critically involved in lexical-conceptual retrieval as with iAG. However, our results suggest that for a word fluency task involving a relatively broad category (animals), the pIFG is not significantly implicated, and less so than the iAG. By contrast, some LSM studies of semantic errors in picture naming implicate pIFG damage, among other regions (Dell et al., 2013; Schwartz et al., 2009). However, picture naming, unlike the word fluency task, involves selecting among competing alternatives (e.g., to name a picture of a cat, the competing alternative *dog* must be suppressed). Interestingly, a recent LSM study of word-level semantic errors in natural, connected speech found that increased errors were associated with temporal and inferior parietal damage, but not frontal damage (Stark et al., 2019). Connected speech differs from confrontation or picture naming in allowing the subject to select alternative words or concepts, which reduces the burden of the task on selection abilities. The whole body of evidence is consistent with a role for pIFG in a selection or control mechanism that is critically involved when there is competition among items, but speaks against a role for basic retrieval of lexical items or associated concepts in frontal cortex, supported instead by temporal and inferior parietal cortex (Lambon Ralph, 2017; Lau et al., 2008; Novick et al., 2005). This selection mechanism could be domain-general (Novick et al., 2005), but subregions of pIFG that respond selectively to language (Fedorenko, Duncan, et al., 2012) could also implement a language-specific control mechanism (Matchin, 2018).

An interesting question is why the aSTS, and the ATL more broadly, was not implicated in semantic category word fluency deficits. Both the aSTS and the iAG have been implicated in semantic processing, broadly construed, in both neuroimaging studies and lesion-symptom mapping of semantic word-naming errors (Binder et al., 2009; Fridriksson et al., 2018; Schwartz et al., 2011). Most pointedly, the degenerative syndrome known as primary progressive aphasia of the semantic subtype (PPA-S, also known as semantic dementia) is strongly associated with mostly left, but sometimes bilateral, atrophy of the ATL, with increasingly stronger deficits in conceptual knowledge and single word comprehension (Hodges et al., 1992; Hodges & Patterson, 2007; Mesulam et al., 2013, 2015). Category fluency has also been shown to be reduced in patients of this type (Hodges & Patterson, 1992). However, our study, as well as Baldo et al. (2006), failed to identify any hint of an effect in the aSTS or the ATL more broadly for semantic category word fluency deficits.

We suggest here that the relevant distinction is concept specificity. In the study reported by Hodges & Patterson (1992), the PPA-S patients had reduced category fluency for more specific categories, breeds of dog and boats, requiring finer differentiation of features than animals, the broader category used here and in Baldo et al. (2006). In fact, Hodges & Patterson (1992) showed that in a picture sorting test, PPA-S patients were able to perform broad categorizations such as living vs. man-made quite well, in stark contrast to their picture naming and word comprehension abilities. Both picture naming and most word comprehension tasks (with picture pointing as the response) require a subject to process specific visual features and attributes of an object. The involvement of the ATL in semantic processing likely relates to specific attributes or features, particularly highlighted in certain word-level production and comprehension tasks, rather than more general ones (Rogers et al., 2006). This is supported by magnetoencephalography studies which have showed that activation in ATL is contingent on concept specificity, e.g. greater activation for higher-specificity words like *canoe* relative to lower-specificity words like *boat*, and that combinatory effects in this area only emerge for lower-specificity words (Westerlund & Pylkkänen, 2014; Zhang, 2015; Ziegler & Pylkkänen, 2016). Concept specificity may also help to explain the purported involvement of the ATL in syntax, as the increased activation in ATL for more complex structures may reflect the increasing concept specificity correlated with structural complexity.

Overall, a picture has emerged by which the ATL is involved in retrieving the features of specific entities, whereas the iAG is involved in a broader semantic function, perhaps involving event representations (Binder & Desai, 2011; Lewis et al., 2015; Matchin et al., 2019; Schwartz et al., 2011). By contrast, the semantic category word fluency task in the present study used a very broad category, animals, which allowed for potentially a wide array of answers without requiring the subject to discriminate highly similar concepts from each other. Thus, performance on this task seemingly critically required iAG but not the aSTS. We would expect that future LSM studies of word fluency using more specific semantic categories, such as dogs, boats, etc., will reveal effects in the aSTS, and possibly the pIFG as well via a selection mechanism.

### 4.4 Word comprehension

Even though the words > nonwords contrast in fMRI often shows activation throughout the language network, including the pIFG (Fedorenko et al., 2016; Fedorenko, Nieto-Castañon, et al., 2012; Matchin et al., 2017, 2019), and the pIFG is reliably implicated in meta-analyses of semantic processing (Binder et al., 2009; Hodgson et al., 2021), our lesion-symptom mapping analyses revealed no significant negative association between damage to the pIFG and behavioral scores on the auditory word comprehension subtest of the WAB-R. Rather, damage was significantly associated with all three of the other selected ROIs (iAG, pMTG, and aSTS, in descending order of significance). Our targeted region by condition interaction analysis showed a robust interaction effect, such that damage to the iAG was significantly more associated with deficits on the word comprehension measure than damage to the pIFG. This converges with the findings from the semantic category word fluency task regarding a dissociation of function between iAG and pIFG, whereby the iAG is critically involved in basic conceptual-semantic retrieval and the pIFG is not. This is consistent with models whereby the iAG plays a role in conceptual-semantic processing that equally supports both production and comprehension (Binder & Desai, 2011; Friederici, 2017; Hagoort, 2016; Lau et al., 2008; Matchin & Hickok, 2020).

### 4.5 Pitfalls of the search for functional selectivity in the language network

Recent neuroimaging studies have shown syntactic and lexical effects distributed across regions of the language network (I. Blank et al., 2016; Fedorenko et al., 2020). Fedorenko et al. (2020) argue that this constitues evidence against the existence of brain areas that selectively process syntax, and evidence for a holistic linguistic architecture in which the lexicon, syntax, combinatorial semantics, and conceptual representations are all intertwined.

First, we strongly caution against using findings from neuroimaging techniques that capture very limited facets of neural structure and function, to inform linguistic architecture, when it is extremely unclear how the postulates of linguistic theory line up with neuroscience (Embick & Poeppel, 2015; Poeppel, 2012; Poeppel & Embick, 2005). Even if it were true that there is no brain region selectively engaged in syntax, or any evidence of functional distinction across regions of the language network, it does not follow that there is no independent syntactic mechanism. The coarseness of the methodologies of fMRI and lesion-symptom mapping do not allow access to many aspects of neuronal function, and a basic syntactic or combinatory mechanism could very well be implemented in subtler biological properties than dedicated chunks of cortical tissue containing thousands of neurons (N. Ding et al., 2016; Gallistel & King, 2010; Matchin & Hickok, 2020; Murphy, 2015).

However, the region by measure interactions we present here, in conjunction with a large historical body of research in aphasiology, lesion-symptom mapping, and functional neuroimaging, suggest that there must be at least *some* differentiation of function within the language network. The key question is what exactly the relevant distinctions are. For example, our results do not imply that the pMTG or pIFG are *selectively* engaged in syntax to the exclusion of the lexicon, or that the iAG is selectively involved in lexical-conceptual retrieval to the exclusion of combinatory processing of any kind. In fact, the quest for evidence of functional selectivity misses what we perceive to be the goals of the neurobiology of language: to identify the functional organization of language in the brain, regardless of the issue of specificity.

In previous work, two of us (Matchin & Hickok, 2020) have suggested that the pMTG and pIFG implement lexical-syntactic functions, with the pMTG processing hierarchical relations stored on individual lexical items and the pIFG processing linear morpho-syntactic relations. We attributed conceptual-semantic processing to different regions, namely the iAG and aSTS. The fact that all of regions in neuroimaging studies respond to lexical experimental manipulations does not speak against these hypotheses. Rather, a contrast such as words > pseudowords is likely to tax multiple functions. For example, the words > pseudowords contrast engages lexical-syntactic processing mechanisms, that is access to the stored repository of words with their associated syntactic frames, access to the meanings associated with individual lexical items, and combinatory semantics enabled by the presence of real words. The fact that all regions of the language network respond to a lexicality manipulation is therefore unsurprising, because this experimental contrast likely engages all of these functions.

Both functional neuroimaging and LSM provide opportunities to uncover the functional architecture of the language network. The evidence we presented here bolsters existing studies by revealing region by measure interaction effects that provides strong evidence of functional dissociations across regions (Nieuwenhuis et al., 2011). Future studies aiming to identify further functional dissociations should develop subtler experimental measures beyond relatively course measures such as comprehension of sentences vs. word lists that are capable of distinguishing among possible underlying functions and should test region by measure interactions if possible.

### 4.6 Limitations of the present work

The biggest limitation of the present work is that while two region by measure interaction effects, syntactic comprehension (p = 0.003) and word comprehension (p = 0.004), were significant, the other two effects, semantic category word fluency and expressive agrammatism, only approached significance, not surviving the correction for multiple comparisons (p = 0.024, p = 0.038). Therefore, these effects should be confirmed by future research. In addition, while LSM is a useful complement to functional neuroimaging, it would help greatly to design fMRI studies carefully enough to obtain significant region by condition interaction effects that can complement the interaction effects obtained here.

Secondarily, while we believe that our measures reflect key aspects of linguistic processing, they can be improved upon and additional research should further develop and refine assessments of different aspects of linguistic processing in people with aphasia. The semantic category word fluency measure was derived from more general test batteries that were not designed to focus on conceptual-semantic processing. Although the existing literature on this measure indicates a strong language component, and a minimal executive function component, it is likely there was at least some executive function contribution to this task. Future research should develop newer, better targeted measures of conceptual-semantic processing that do not involve selection among competing alternatives (as in picture naming) or executive function demands (as in the Pyramids and Palm Trees test). The expressive agrammatism measure was a perceptual rating by experts. Casilio et al. (2019) showed strong concurrent validity between quantitative measures of agrammatism and perceptual ratings of grammatical speech deficits. Our own perceptual ratings of agrammatism had very high inter-rater reliability (Matchin et al., 2020; Den Ouden et al., 2019). Thus, we believe our perceptual ratings are justified. However, it would be useful to complement these perceptual ratings with objective, quantifiable measures that would complement these results. The syntactic comprehension measure, standard in the literature on syntactic comprehension, might involve additional processes beyond syntax such as combinatorial semantics or working memory. While the region by measure interaction effect we showed here indicates a functional dissociation between the function of pMTG and pIFG, it does not necessarily provide knock-down evidence for a strictly syntactic function of pMTG. Additional measures that seek to isolate syntax from other mechanisms would help to clarify the picture.

Finally, one potential objection to our conclusions regards the possibility of post-stroke functional reorganization. If language is reorganized in the brains of those suffering a stroke to language-relevant regions, as suggested by some authors (Hartwigsen & Saur, 2019; Stefaniak et al., 2020; Turkeltaub, 2019), how does this impact our conclusions? First, to the extent that there is functional organization in post-stroke aphasia, it is likely facilitatory rather than fundamental (e.g. Fridriksson et al., 2012). People with chronic post-stroke aphasia retain significant deficits, and a recent meta-analysis and review of functional neuroimaging studies in people with post-stroke aphasia suggesting no evidence of large-scale reorganization (Wilson & Schneck, 2020). Secondly, functional reorganization would only weaken our ability to detect distinctions across regions, as the impact of a lesion on a given function would be weakened by recovery of function. However, lesion-symptom mapping studies readily identify strong correlations between linguistic deficits and patterns of brain damage, and these findings correspond to what is known from functional neuroimaging (Fridriksson et al., 2018). Crucially, lesion methods are an essential complement to functional neuroimaging in that it helps to identify causal mechanisms of cognition in the brain, both in human and non-human organisms (Lomber et al., 2010; Milner & Goodale, 1995; Rorden & Karnath, 2004). Theoretical models of language and the brain were originally developed from the study of aphasia (Wernicke, 1874) and have continued to incorporate its insights (Bornkessel-Schlesewsky & Schlesewsky, 2013; Friederici, 2002; Hagoort, 2005; Hickok & Poeppel, 2000, 2004, 2007; Pinker & Ullman, 2002; Rauschecker & Scott, 2009). This impact will only be strengthened with improved methods and sample sizes.

## Funding & Acknowledgments

This research was supported by National Institute on Deafness and Other Communication Disorders grants P50 U01 DC011739 and R01 DC014664 awarded to Julius Fridriksson, and a National Institute on Deafness and Other Communication Disorders grant awarded to Alexandra Basilakos, T32 DC014435. We would like to thank Grigori Yourganov and Jonathan Venezia for advice on statistical analysis. We would also like to thank Leigh Ann Spell, Allison Croxton, Anna Doyle, Michele Martin, Katie Murphy, and Sara Sayers for their assistance with data collection, and graduate student clinicians in the Aphasia Lab for transcribing and coding speech samples.

